# Multidimensional stimulus-response correlation reveals supramodal neural responses to naturalistic stimuli

**DOI:** 10.1101/077230

**Authors:** Jacek P. Dmochowski, Jason Ki, Paul DeGuzman, Paul Sajda, Lucas C. Parra

## Abstract

In neuroscience, stimulus-response relationships have traditionally been analyzed using either encoding or decoding models. Here we combined both techniques by decomposing neural activity into multiple components, each representing a portion of the stimulus. We tested this hybrid approach on encephalographic responses to auditory and audiovisual narratives identically experienced across subjects, as well as uniquely experienced video game play. The highest stimulus-response correlations (SRC) were detected for dynamic visual features. During narratives both auditory and visual SRC were modulated by attention and tracked correlations between subjects. During video game play, SRC was modulated by task difficulty and attentional state. Importantly, the strongest component extracted for visual and auditory features had nearly identical spatial distributions, suggesting that the predominant encephalographic response to naturalistic stimuli is supramodal. The variety of novel findings demonstrates the utility of measuring multidimensional stimulus-response correlations.

## Introduction

Understanding the relationship between a sensory stimulus and the resulting neural response is a fundamental goal of neuroscience. Two distinct paradigms have shaped the pursuit of the neural code. The *encoding*approach attempts to explain neural responses from features of the stimulus, typically via linear filtering [7]. Examples include receptive fields and spike-triggered averages in single-unit electrophysiology [7], the generalized linear model (GLM) in functional magnetic resonance imaging (fMRI) [14, 45], spectrotemporal response functions (STRF) in electrocorticograms [10], and temporal response functions in encephalographic recordings [38, 37, 8]. In contrast to encoding, the *decoding*approach is to predict the stimulus by filtering over an array of neural responses. Decoding techniques have been shown to reconstruct sensory experience in a large number of findings spanning animal [3, 68, 62, 5] and human investigations of both visual [49, 65, 44, 32, 48, 26] and auditory stimuli [53, 41, 43, 42, 51].

The encoding and decoding approaches possess opposing strengths and weaknesses: whereas encoding models operate on the stimulus and are thus easily interpretable [46], they generally predict the responses of individual data channels (i.e, neurons, voxels, or electrodes) and do not efficiently recover distributed neural representations. Decoding techniques filter neural activity over multiple channels and are therefore naturally suited to capturing distributed representations, but at the expense of models that are often difficult to interpret and prone to overfitting. Therefore, an approach that efficiently captures distributed neural representations and is readily interpretable in the stimulus space is lacking.

Here we propose a hybrid approach that combines the strengths of encoding and decoding. The technique integrates neural responses across space while filtering the stimulus in time, i.e. it “decodes” neural activity to recover an “encoded” version of the stimulus. By jointly learning decoding and encoding models, distributed neural representations are identified and explicitly linked to portions of the stimulus. In contrast to existing paradigms, this approach decomposes the neural representation of stimuli into multiple dimensions, with each dimension defined by a (spatial) response component and a (temporal) stimulus component.

We evaluated the technique on recordings of neural activity in response to various naturalistic audiovisual stimuli and found multiple significant dimensions of stimulus-response correlation (SRC) for auditory and visual features. These multiple dimensions were variably modulated by the attentional state of the observer. Interestingly, we found that independent visual and auditory features possessed a common response component, suggesting that the dominant neural response to natural stimuli is supramodal.

Inter-subject correlations (ISC) of neural responses to natural stimuli have recently been shown to reflect a variety of behaviors [19, 22, 35, 67, 63, 12, 11]. We expected that stimulus responses would be similar across subjects and thus predicted that SRC would track ISC; indeed, SRC covaried with ISC for both auditory and visual features. In contrast to ISC and event-related potentials, however, the proposed method does not require repeated exposures to the stimulus and is thus applicable to the study of unique experience. We therefore studied neural activity during video game play and identified SRCs that reflected both the game difficulty and attentional state of the player. The variety of novel findings attests to the utility of the hybrid encoding-decoding approach.

## Results

The hybrid encoding-decoding technique is developed in the Methods section, with details of implementation provided as Supplementary Information (SI). Briefly, encephalographic activity is spatially filtered (decoded), whereas stimulus features are temporally filtered (encoded). The weights of these spatial and temporal filters are selected to maximize the correlation between the neural response and stimulus: the *stimulus-response correlation (SRC)*. This optimization problem is efficiently solved by canonical correlation analysis (CCA), which provides a set of filters that decompose the neural response into *response components* and the stimulus into *stimulus components*. Each pair of components captures independent dimensions of the SRC. In the following we apply the technique to naturalistic audiovisual stimuli in order to investigate the brain’s representation of multiple dynamic stimulus features and their dependence on attentional and task demands.

### Dynamic visual features elicit strong multidimensional SRC

We first sought to determine which features of naturalistic stimuli evoke the strongest SRC. For a popular film clip during which we recorded the evoked scalp potentials of *N* = 30 viewers, we extracted a set of visual and auditory features including optical flow, visual temporal contrast, sound amplitude envelope, luminance, and spatial contrast (see Methods). Applying these derived features and neural responses to the hybrid encoding-decoding scheme led to a range of SRC values (Fig. 1A). We found statistically significant correlations along multiple dimensions (i.e., component pairs) for four of the five features (all *p* < 0.05, computed using phase-randomized surrogate data). SRCs of different component pairs are shown cumulatively, as each pair captures a different dimension in the data with uncorrelated activity. The strongest SRCs were exhibited by temporal contrast and optical flow, exceeding the correlations with sound envelope (paired t-test for sound envelope with temporal contrast, *t*(29) = 5.7, *p* = 4 × 10^*−*6^, and with optical flow *t*(29) = 4.7, *p* = 5 × 10^*−*5^).

**Figure 1:**
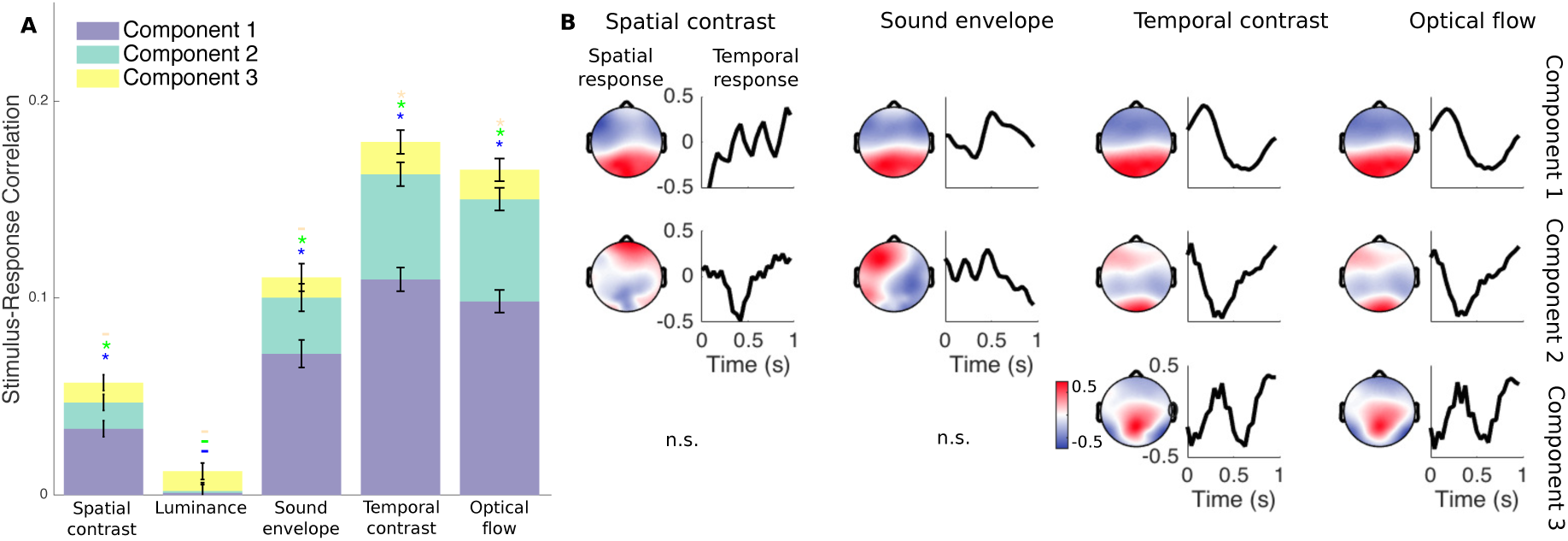
Neural responses to video track dynamic visual features. **A** SRC as computed for various auditory and visual features of a clip from “Dog Day Afternoon”. Significant correlations were detected along multiple dimensions for 4 of the 5 features considered (*p* < 0.05, and indicated by ‘*’ with color indicating the component tested). Correlations with temporal contrast and optical flow, features that differentiated pixel values across frames, exceeded those with the sound envelope (*p* < 0.04, paired t-test). **B** Spatial responses convey the topography of neural activity that best expressed the stimulus features. Temporal responses reflect the delays between stimulus and neural response. While the topographies of the first response components (left panels in top row) were congruent for all features, the temporal responses were not. This suggests a “multiplexing” of multiple stimulus features onto a common neural substrate.

By construction, the response components recovered by CCA are temporally uncorrelated with one another. However, when regularizing covariance as was performed here (see SI 2), the response components may exhibit some level of correlation. Thus, to confirm that regularization did not introduce “cross-talk”, we also measured the correlation between mismatched stimulus and response components (e.g. the correlation between stimulus component 1 and response component 2). These correlations were found to be very low (mean ± standard deviation across all features and component pairs: 0.0005 ± 0.003). Comparing this to the SRC measured within matched component pairs (as high as 0.1 for temporal contrast), it is clear that the multiple response components detected by the hybrid technique were distinct.

### Multidimensional SRC reveals supramodal component

Each SRC was formed by extracting a response component that was correlated with a stimulus component. Response components possessed a topography, termed a *spatial response*, that captured the spatial distribution of the recovered neural activity. Stimulus components were extracted by a temporal filter that captured a *temporal response*. Together, the spatial and temporal responses fully convey the mapping between original stimulus and evoked neural response (see SI 3 for details). Interestingly, the spatial responses of the first component were similar for all features (Fig. 1B). On the other hand, the associated temporal responses differed across features, and often exhibited oscillations. This suggests that multiple stimulus features are represented by a common neural substrate that is selective for different frequencies of naturalistic stimuli.

While the finding of congruent spatial responses was expected for the two dynamic visual features which were strongly correlated (correlation between optical flow and temporal contrast: *r* = 0.96), we did not expect to find such similar topographies for weakly correlated auditory and visual features (correlation between sound envelope and temporal contrast: *r* = 0.067). To rule out that the similarity of the auditory and visual spatial responses was due to this small correlation, we subtracted from the temporal contrast the fraction that was explained by the sound envelope, and vice versa. In doing so we generated orthogonal time series for temporal contrast and sound envelope. The spatial responses of the first component for these uncorrelated visual and auditory features still had nearly identical distributions on the scalp (Figure 2, correlation between spatial responses of sound envelope and temporal contrast: *r* = 0.99). We interpret this as the spatial filter picking up the activity of a supramodal neural circuit that dominates the EEG evoked response. Note that the spatial filter weights themselves were also quite similar for the auditory and visual features (correlation between spatial filter weights: *r* = 0.88). The associated temporal filters, which reflect the time delays with which the stimulus evokes a response, were inversely related (correlation between temporal filter weights: *r* = −0.83). This indicates that auditory and visual features drove the supramodal response with opposing polarity in the EEG.

**Figure 2:**
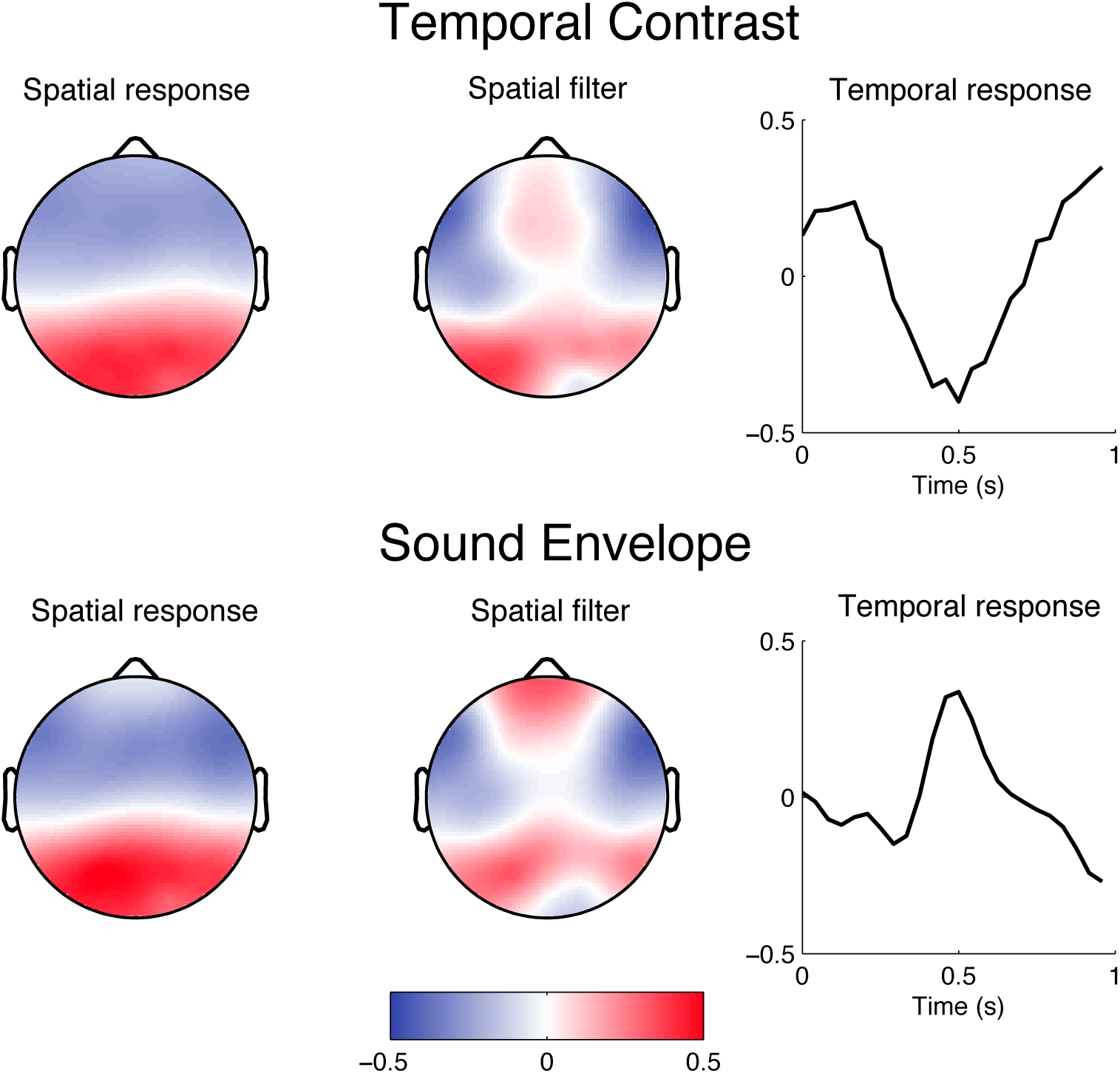
Auditory and visual features evoke congruent spatial responses. The spatial responses of the first component for visual temporal contrast and sound envelope, which depict where on the scalp these stimuli were expressed, were highly similar (left column, *r* = 0.99), even though the features were decorrelated (*r* = 0). The spatial filters that extracted the response components were also highly correlated (middle column, *r* = 0.88). However, the associated temporal responses were inversely related (right column, *r* = −0.83). This suggests that a common neural substrate is driven in a supramodal fashion, but with opposite sign for visual and auditory features. All values have arbitrary units as SRC is independent of scale.

### Multidimensional SRC tracks Inter-Subject Correlation (ISC)

A number of reports show that dynamic natural stimuli elicit similar responses across subjects in fMRI, EEG and MEG [19, 11, 39]. For responses to be reproducible across subjects they also have to be evoked reliably. Therefore, we hypothesized that there would be a correspondence between how strongly the stimulus drove individual neural responses and how reliable the responses were across subjects. To test this, we computed a *time-resolved* measure of the SRC (by summing across the first three component pairs of the hybrid technique) for the temporal contrast and sound envelope of the same film clip. Similarly, we also measured the time-resolved ISC (by summing across the three components maximizing correlation across subjects) experienced during the same stimulus (see Methods). In line with our hypothesis, a significant portion of the variability in the ISC time series could be explained from both visual and auditory SRC (Fig 3; temporal contrast: *r* = 0.59, *p* = 3 × 10^*−*31^; sound envelope: *r* = 0.34, *p* = 3 × 10^*−*10^). This suggests that the exogenous drive provided by the common stimulus underlies the reliability of neural responses across subjects.

**Figure 3:**
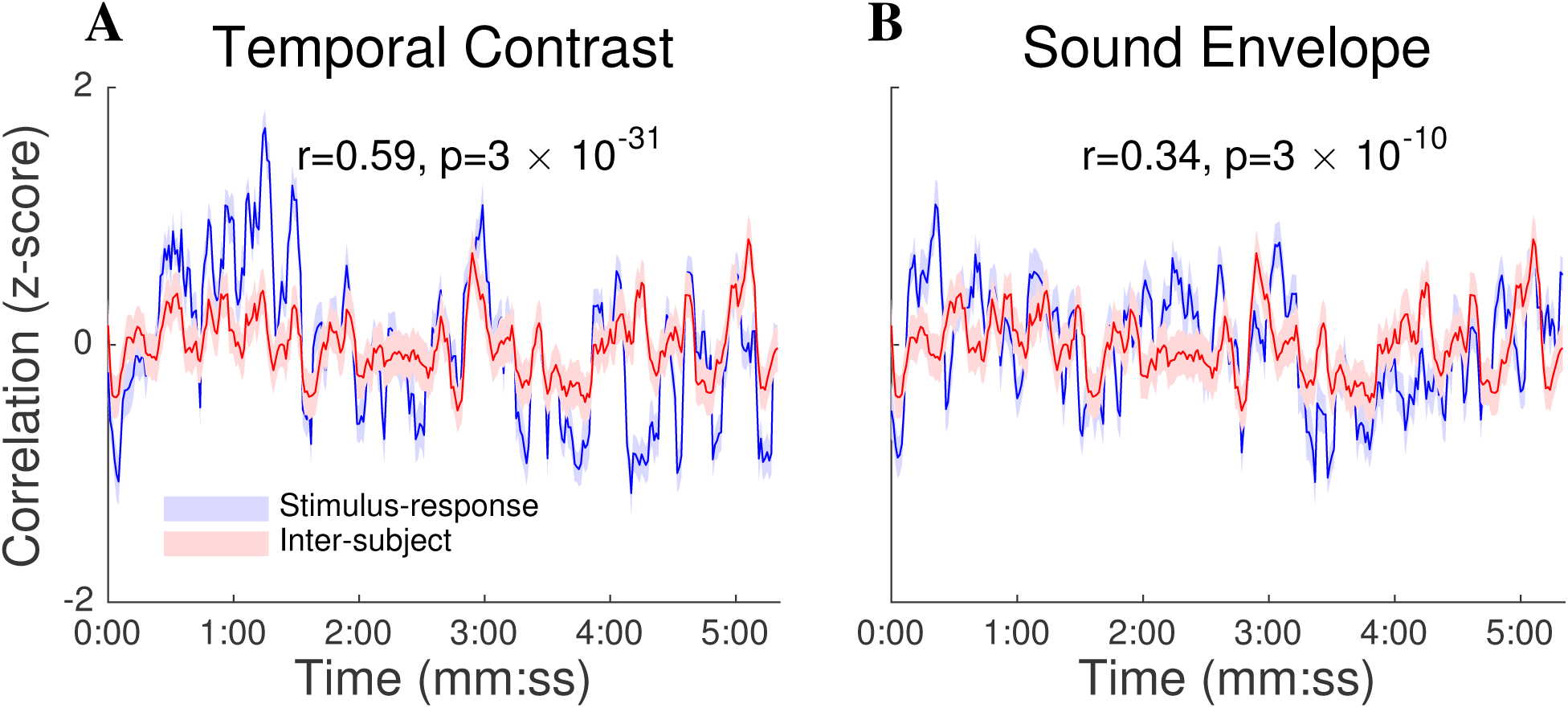
SRC tracks the inter-subject correlation (ISC) of neural responses to naturalistic stimuli. **A** The time course of the SRC, as computed on the temporal contrast of a film clip, explains 34% of the variability in the ISC time course (*r* = 0.59, *p* = 3 × 10^*−*31^). This suggests that the exogenous drive provided by the common stimulus underlies the reliability of neural responses across subjects as measured by the ISC. **B** Same as (A) but now for the envelope of the film’s soundtrack. The SRC accounts for approximately 12% of the variability in the ISC (*r* = 0.34, *p* = 3 × 10^*−*10^). Shading indicates the SEM across subjects (for SRC) or subject pairs (for ISC).

### Multidimensional SRC is modulated by attentional state

Given the long-standing evidence showing that attention modulates evoked responses (e.g. [55]), we hypothesized that the SRC would decrease when the level of attention directed to the stimulus was reduced. To test this, we reanalyzed previous data recorded in two attentional conditions [33]: subjects either naturally attended to the stimulus or performed a counting task, intended to distract viewers from the stimulus. Two film clips (“Bang! You’re Dead” and “The Good, The Bad, and the Ugly”) and one narrated story (“Pie Man”) were considered for this analysis. All subjects were presented the same stimuli under both conditions (i.e., we took repeated measures). We investigated whether SRC was modulated by attention (attend vs count) and if this was specific to particular stimuli or component pairs. We first analyzed the SRC using the sound envelope. A three-way, repeated-measures ANOVA with component, attention and stimulus as factors identified main effects of attention (*F* (1) = 7.48, *p* = 0.008) and stimulus (*F* (1) = 29.17, *p* = 1.82 × 10^*−*9^) and an interaction between component and stimulus (*F* (2) = 14.15, *p* = 1.02 × 10^*−*5^). Follow-up pairwise comparisons showed that the reduction in SRC for the “count” condition was driven by the two stimuli containing speech (see Figure 4A). In contrast, the “The Good, the Bad, and the Ugly” had minimal speech content and elicited weak SRC that was not modulated by attention. This suggests that the effect of attention on auditory EEG responses may be specific to speech. For the two audiovisual clips, we measured SRC using the optical flow (Fig 4A) and again performed a three-way, repeated-measures ANOVA with attention, stimulus and component as factors. There was again a strong main effect of attention (*F* (1) = 34.6, *p* = 10^*−*5^), with reduced SRC in the “count” condition. An interaction between attention and stimulus component (*F* (1) = 12.2, *p* = 0.002) was also found. Specifically, “Bang! You’re Dead”, which has a more suspenseful narrative was more robustly modulated by attention, consistent with results reported in [33].

**Figure 4:**
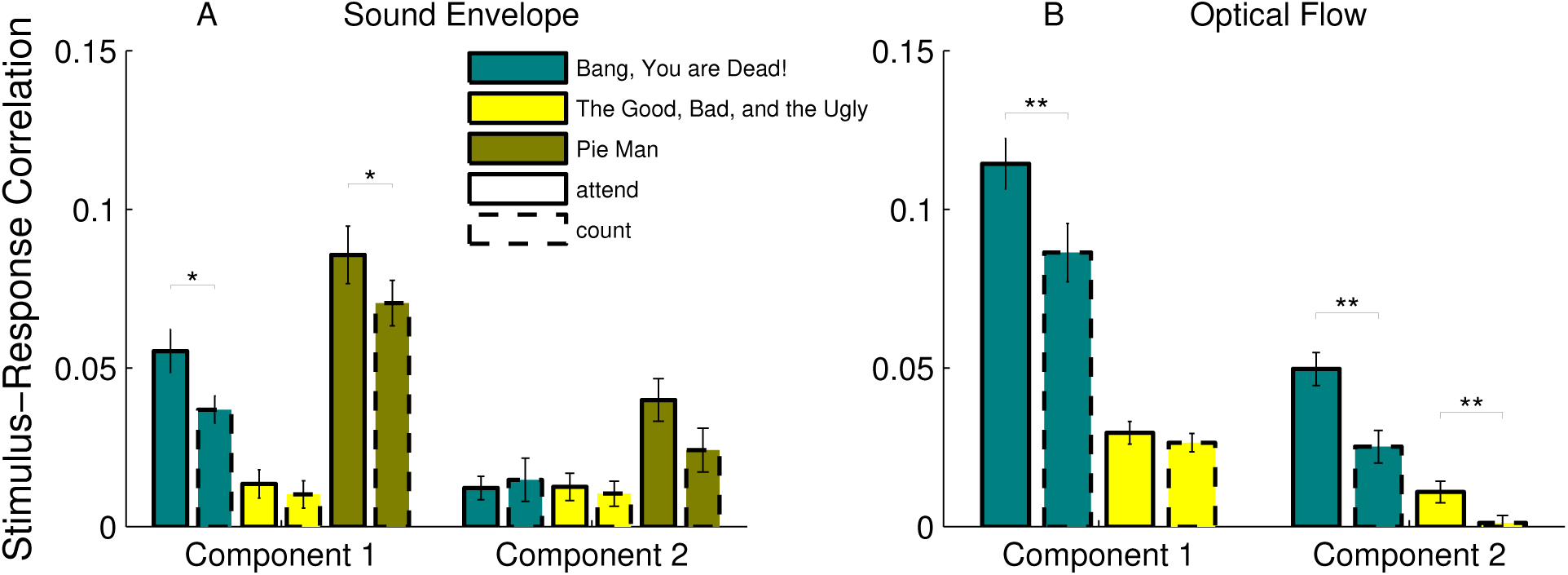
Multidimensional SRC is modulated by the level of attention directed to the stimulus. Subjects viewed film clips or listened to an audiobook while either naturally attending to the stimulus (‘attend’) or performing a counting task (‘count’). **A** SRC as measured for the sound envelope of two audiovisual and one auditory stimulus. A comparison of SRC between count and attend conditions showed a significant effect of attention (see ANOVA resuls in text). Follow-up comparisons indicated that SRC was modulated by attention in the first component but only for the stimuli that contained speech (t-test, BYD: *p* = 0.021 and PM: *p* = 0.044). **B** SRC with optical flow was weaker for the count condition as compared to the attend condition (see ANOVA results in text). Follow-up comparisons indicated that this effect was robust in the first component for BYD (t-test: *p* = 0.002), and in the second component for both audiovisual stimuli (t-test, BYD: *p* = 0.0011 and GBU: *p* = 0.0036)

### Hybrid technique captures uniquely experienced SRC

To demonstrate that the proposed technique can capture SRC elicited by uniquely experienced stimuli, we recorded scalp potentials from *N* = 5 subjects while playing a car-racing video game (Fig 5A). The ongoing feedback between player and game meant that every race was perceptually unique. After reconstructing the optical flow of the video game display during each race, the stimulus time series and neural responses were used to measure SRC with the hybrid technique. The first response component possessed a left-lateralized spatial response focused over the parietal electrodes (Fig 5B, left panel), but possibly also consistent with right-hand motor activity. This component had an early response (i.e., 150 ms) to the game’s optical flow (Fig 5B, temporal response in the right panel). Interestingly, the first stimulus and response components of optical flow derived during the video game differed from those extracted during passive film viewing (compare Fig 5B with Fig 1B). This discrepancy may indicate that when actively engaged with natural stimuli, distinct neural circuits are recruited. An alternative interpretation is that somatosensory and motor activity correlated with optical flow as players controlled speed and direction with right-hand key-presses. In this case, however, one would have expected a more central or anterior spatial response.

**Figure 5:**
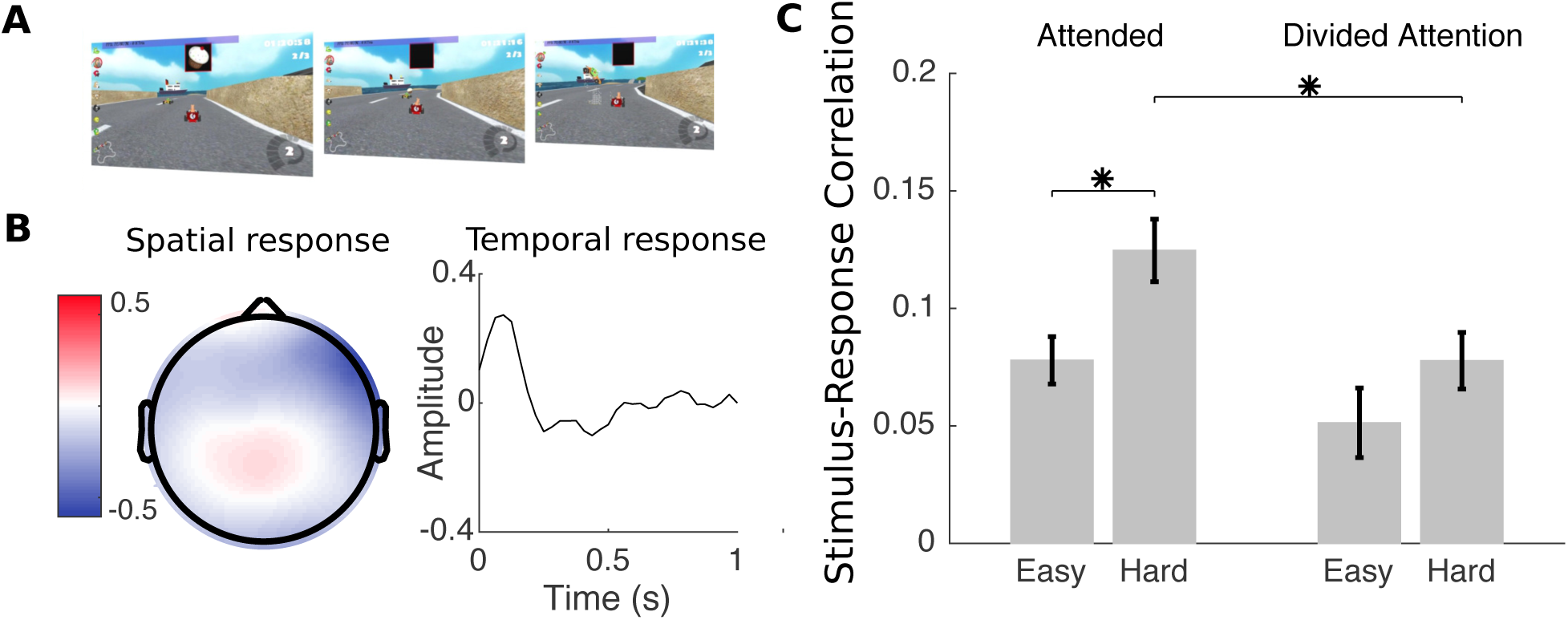
SRC with optical flow measured during video game play is modulated by task difficulty and attention to a secondary task. **A** Neural activity was recorded during play of a racing video game, and the optical flow of the interactive stimulus was computed. **B** The spatial response of the component that best tracked the game’s optical flow was focused at parietal electrodes with a left lateralization. The associated temporal response peaked early at 150 ms. **C** Dominant correlations (first component pair) increased during difficult runs (t-test, *p* = 0.03), while *decreasing* when the player’s attention was divided between the game and a secondary task (t-test, *p* = 0.03).

### SRC is modulated by difficulty and presence of dual-task

The video game consisted of a car race with obstacles and competing drivers. Players experienced two levels of race difficulty by varying the number and skill of competing drivers. The game also included a divided-attention condition which required players to simultaneously attend to the top center of the screen (Fig 5A), where items were presented for selection [15]. We predicted that players would devote more resources to the driving task during difficult races, resulting in higher SRC. In contrast, during periods of divided attention to a secondary task, SRC would be reduced. Two-way, repeated measures ANOVA on SRC with optical flow revealed a main effect of both difficulty (*F* (1) = 8.35, *p* = 0.01) and attention (*F* (1) = 8.42, *p* = 0.01), with correlations increasing during difficult races and decreasing during the divided attention task, as predicted. Follow-up tests confirmed the presence of significantly larger correlations during difficult races in the attend condition (paired t-test, *p* = 0.03), and significantly lower correlations during divided-attention races on the hard condition (paired t-test, *p* = 0.03). It is important to note that the stimuli differed with game difficulty, and thus one cannot rule out that the effects are a result of varying stimuli and not of varying neural responses.

### Hybrid technique extracts stronger SRC than encoding or decoding alone

Finally, we sought to compare the magnitude of SRC detected by the hybrid technique with that measured by existing approaches. Since the hybrid encoding-decoding technique filters both the stimulus and the neural response before correlating them, we expected that this technique would yield stronger correlations than those obtained by the encoding or decoding approaches alone (see Discussion). For all features that elicited significant SRC (see Fig 1), we applied an established encoding technique [38, 37] to measure the correlation between the evoked EEG responses and those predicted from the stimulus features. Similarly, we also measured the correlation between the stimulus feature and that estimated from the neural activity using a common decoding technique [53]. We then contrasted these values to the multdimensional SRC computed by the hybrid technique. When accounting for the SRC of all significant component pairs, the hybrid approach outperformed both encoding and decoding for all features (paired t-test, *p <* 8 × 10^*−*8^). When only considering the first SRC dimension, the proposed technique outperformed encoding for all features (paired t-test, *p* < 0.04), but decoding for only two of four features (paired t-test, *p* < 6.9 *×* 10^*−*5^).

## Discussion

Here we have developed a hybrid technique for learning the mapping between a dynamic stimulus and the corresponding neural response. By simultaneously encoding the stimulus and decoding the neural response, the proposed approach recovers multiple dimensions of SRC via the canonical correlation analysis formalism. We found that visual and auditory features drove a common neural substrate in a supramodal fashion, even after removing all correlation between the film’s soundtrack and visual features. Moreover, the dimensions of the SRC were variably modulated by attention. The SRC was also shown to closely track the across-subject correlations (i.e., ISC) of neural responses that have recently been employed to decode a variety of cognitive states. In contrast to the ISC, however, the multidimensional SRC does not require repeated exposures to the stimulus. The technique is thus applicable to the study of unique experiences, as was demonstrated here for video game play, where both attentional and task demands were shown to modulate the SRC.

### EEG tracks dynamic visual features

While there have been multiple reports of EEG responses tracking auditory features, in particular the envelope [37, 10, 9, 57, 16, 8], relatively little comparable findings exist for visual features (see [18] for an exception). The hybrid technique developed here demonstrated that dynamic visual features were correlated with encephalographic responses to a level comparable if not stronger than the well-studied auditory envelope.

### Supramodal component

The supramodal component identified here bears a strong resemblance to the component that was previously found to maximize the ISC of EEG responses to film clips and advertisements [12, 11, 33]. In these earlier studies, the supramodality was obscured as the ISC approach is blind to which features of the stimulus drive reliable responses. Here we demonstrated that both auditory and visual features correlated with this component. It is thus possible that this component is selective to integrated audiovisual activity, as has been observed in temporal cortex during presentation of speech [6] as well as individual letters [58]. Alternatively, the activity may be related to attentional networks that are entrained by the stimulus regardless of modality [36, 66]. Due to the inherent challenges of localizing EEG components [17], we are cautious in relating this supramodal component to a specific cortical origin. Rather, additional studies employing imaging modalities with finer spatial resolution (e.g. fMRI), in conjunction with experimental designs where SRC are measured for both uni-and multi-sensory presentations, are needed.

### Distributed stimulus representations

A basic premise of the proposed approach is that stimulus features are represented by distributed rather than local neural responses. This is particularly true for EEG where the activity from a localized neural population can be detected at multiple electrodes. The encoding approach, conventionally used for analysing spiking activity [7], fMRI [14] and recently also EEG [38], models neural responses at individual channels (electrodes, voxels), and does not directly leverage such distributed activity. Decoding approaches, in contrast, can combine responses that are distributed and appear only weakly in individual channels [31, 49]. While encoding models are sometimes reversed to provide decoding [48], such an approach often ignores the correlated nature of neural responses [13]. The hybrid encoding-decoding technique captures distributed representations as components of the neural response. These components are linked to temporal components of the stimulus, and thus readily interpretable.

### Separating multiple dimensions of SRC

An important aspect of the proposed technique is its ability to extract multiple, independent dimensions of SRC. These multiple dimensions allow one to more finely probe the effects of experimental manipulations. For example, consider the effect of attention on SRC during film viewing (Figure 4). For the sound envelope, only the correlation of the first component was modulated by altering attentional state. Conversely, for optical flow, the modulation was strongest in the space of the second component. Thus, the multidimensional nature of the proposed approach allows one to identify the neural circuits driven by experimental variables.

### Relation to Inter-Subject Correlations

The multidimensional SRC was shown to track the inter-subject correlation (ISC) between neural responses to the same stimulus. This result is significant because unlike the ISC approach, SRC may be measured with only one subject and one exposure to a stimulus. There are several recent examples of the ISC reflecting behavioral outcomes. For instance, ISC predicts subsequent memory of the stimulus [19], is indicative of viewer engagement [12, 11], correlates with the effectiveness of communications between individuals [63, 61], and reveals the time scale of information integration for narratives [22]. With the proposed technique, many of these studies can now potentially be performed in the context of unique experience. This is demonstrated here using video game play, which is adaptive to user behaviour and therefore necessarily results in a different stimulus for every rendition of the game.

### Attentional modulation of evoked activity

Manipulating attentional state has been previously shown to strongly affect the ISC of the encephalogram [33], and preliminary evidence [56] suggests that this may partly result from varying evoked response magnitude, which is known to be affected by attentional state. We thus reanalyzed the data from [33] and found that both auditory and visual SRC were indeed modulated by attention (Figure 4). Modulation of SRC with attentional tasks has been previously demonstrated for the cocktail party problem (i.e., attend to the voice of speaker A vs speaker B when both are simultaneously speaking). Both decoding [42, 52, 16] and encoding approaches [10] have been used in this context. It is interesting that the attentional modulation of the SRC with sound envelope was found here only for stimuli that contained speech, indicating that the modulation was not due to generic sound-evoked responses. We also found a robust modulation of SRC for the visual feature, but this too depended on the specific stimulus. Once again, this indicates that attentional modulation may not be a generic property stimulus evoked responses, and may explain the mixed results found in studies of task-related visual attention [50, 60].

### Interpreting video game activity

We found that the neural response to optical flow differed both spatially and temporally depending on whether the subject was passively observing or actively engaged with the stimulus. During active play of a video game, a response component with a parietal topography and early time course emerged. This was in contrast to the slower supramodal component that was found to best correlate with optical flow during passive film viewing. While it is tempting to speculate that this result is evidence of mode-dependent visual processing, we cannot rule out alternative explanations that involve the effects of motor actions on the EEG: during video game play, subjects continually pressed keyboard buttons to control the game. Even though the observed spatial response is not consistent with a motor topography, there is evidence that button presses alter the task-evoked topographies of oddball paradigms [59]. Further manipulations that control for the effects of key presses will need to be conducted to pin down the source of the response component recovered during video game play.

### Interpreting the magnitude of SRC

When comparing the hybrid technique to existing approaches (Figure 6), we reported the SRC across multiple dimensions. In contrast, traditional encoding or decoding techniques measure SRC in a single dimension, either in the space of the stimulus or response. While for encoding models this can be potentially conducted for multiple channels, it is difficult to ascertain, without additional processing, what fraction of the SRC is shared or unique. With the hybrid approach, the SRC is partitioned into independent dimensions that correspond to different temporal portions of the stimulus. Note that even when considering only the first SRC dimension, the proposed approach often captures more variance in the data. One possible explanation for this is that the hybrid technique filters both the stimulus and response, and can thus remove unexplainable variance prior to computing the SRC. In contrast, when measuring the correlation between the neural response and that predicted by an encoder, there will be significant variance in the actual response that is not stimulus-related. Similarly, when measuring correlation between the stimulus and the output of a decoder, there may be significant parts of the stimulus that are not captured in the neural response. Note also that encoding and decoding techniques apply a separate temporal filter to each channel, thus increasing the number of parameters in the model. The hybrid approach, as developed here, employs a single temporal filter that is shared among the the channels comprising each component, and that may be sufficient to account for the delay between a stimulus and its neural response. With fewer degrees of freedom in the hybrid model, we were generally able to achieve higher cross-validated correlations.

**Figure 6:**
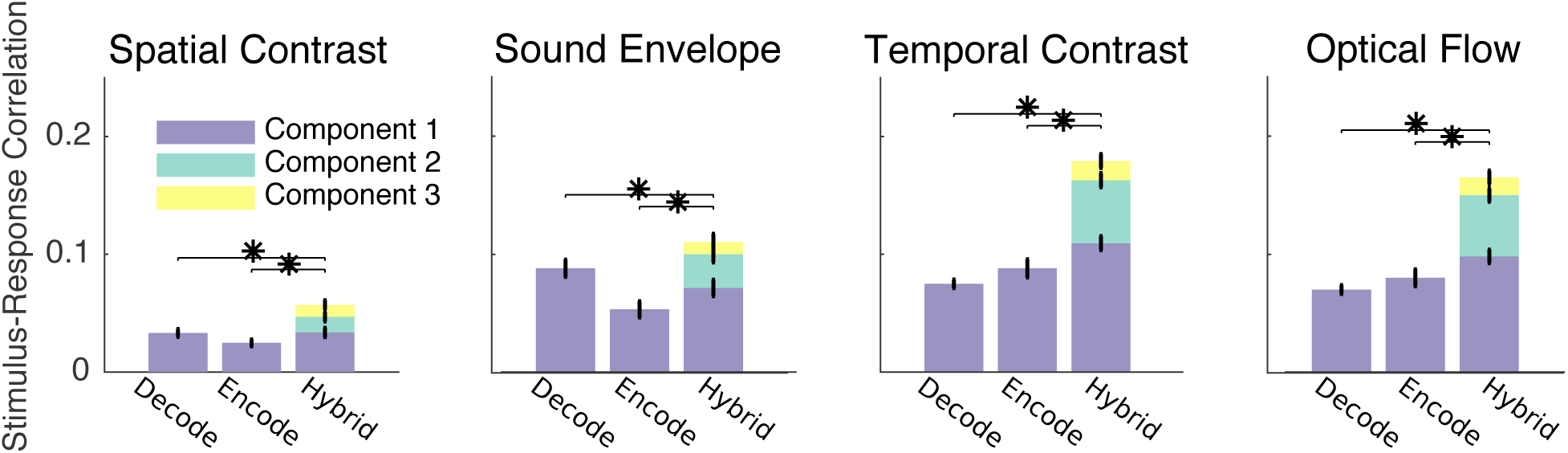
Hybrid encoding-decoding detects stronger SRC than encoding or decoding alone. The SRC captured by established decoding (“decode”) and encoding (“encode”) approaches, as well as the proposed hybrid technique, on encephalographic responses to a film clip. The SRC of the first component pair as recovered by the hybrid technique outperformed encoding or decoding in 6 of 8 comparisons (paired t-test, *p* < 0.04). When accounting for all significant components pairs, hybrid encoding-decoding detected stronger correlations in all cases (paired t-test, *p* < 8 *×* 10^*−*8^).

### Extensions to non-linear architectures and microelectrode arrays

Our ability to uncover the principles of sensory representation goes hand in hand with the ability to explain variance in the neural response. Here the relationship between stimulus and response was constrained to multiple linear mappings. It is expected that the incorporation of more sophisticated architectures that capture non-linear mappings will increase the magnitude of observed SRC. For example, deep neural networks that can synthesize complex functions and account for higher-order correlations may be implemented in a regression. A deep-learning extension of classical CCA has recently been formulated [67, 2]. Kernel methods that exhibit robustness to overfitting may also prove useful [1, 4]. Finally, we note that the formalism presented here is equally applicable to other types of neural data including magnetoencephalography (MEG), fMRI, and multi-unit activity.

## Methods

Here we develop the proposed technique by relating it to the two predominant approaches for the analysis of neural signals: predicting the neural response from the stimulus (encoding), and recovering the stimulus from the neural response (decoding).

### Encoding: modeling the mapping from stimulus to response

Consider a stimulus whose time-varying features are encapsulated by signal *s*(*t*). For an auditory stimulus, the values of *s*(*t*) may represent the sound pressure envelope. For a visual stimulus, *s*(*t*) may represent the luminance. The stimulus is presented to an observer, generating a neural response *r*_*i*_(*t*) in the *i*th data channel (for example, an electrode in a microelectrode or EEG array, or a voxel in fMRI). Encoding seeks to identify the mapping from *s*(*t*) to *r*_*i*_(*t*). This is conventionally performed by filtering *s*(*t*) to produce an estimated neural response 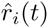 for each channel:

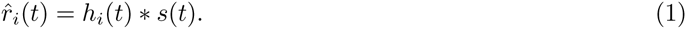

When the response is the firing rate of neurons, the The filters, *h*_*i*_(*t*), could represent the post-stimulus time histogram for each neuron *i*for spiking activity [7]. For EEG the filters represent the evoked response for each electrode *i* [38, 37]. Encoding filters *h*_*i*_(*t*) are found by maximizing the correlation between the observed neural response *r*_*i*_(*t*) and the estimated neural response 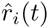 (or a non-linear version of 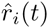 in the case of generalized linear models used in fMRI [14]). Here the notation *∗* denotes temporal convolution, such that 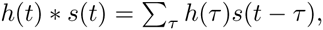 but it could also represent spatial convolution to model visual receptive fields [7]. Note that this optimization problem is generally solved separately for each channel *i* = 1…*D*. Thus, the encoding approach does not leverage potentially distributed representations where the stimulus elicits correlated responses across multiple channels. In particular, if the response in a given channel is below statistical detection, it may be “missed” by an encoding approach.

### Decoding: recovering the stimulus from the response

To remedy this, decoding techniques combine the neural responses of multiple channels and aim to reconstruct the stimulus (e.g. [42, 14]):

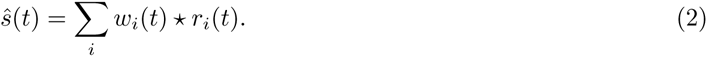

The decoding weights *w*_*i*_(*t*) perform both spatial *and* temporal filtering, and are found by maximizing the correlation between the observed stimulus *s*(*t*) and the estimated stimulus 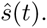. Here the notation ⋆ denotes temporal correlation, such that 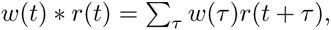 captures the response after stimulus presentation at time *t*. Note that the responses of multiple channels are linearly combined to recover a single stimulus feature, and that the model parameters are now learned jointly. One limitation of this technique is that it is difficult to directly interpret the decoding coefficients *w*_*i*_(*t*) [23]. Moreover, in cases with many channels and long temporal apertures, conventional decoding techniques are prone to over-fitting and thus require careful model regularization [69, 54, 53, 24].

It should be noted that both encoding and decoding techniques can be inverted (to become decoders or encoders) if the unconditional distributions of the stimulus or the response, respectively, are available [46, 48, 47]. While the statistics of the stimulus may be readily estimated, it may be more challenging to estimate the statistics of the response independently of the stimulus.

### Hybrid encoding and decoding

Here we propose a combination of the encoding and decoding approaches by simultaneously filtering the stimulus in time and the neural responses in space:

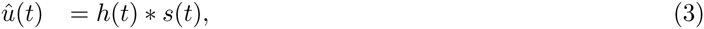

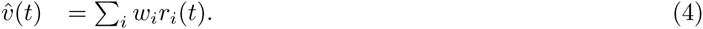

The encoder *h*(*t*) and decoder *w*_*i*_ are found by maximizing the correlation between the encoded stimulus 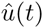 and the decoded response 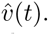 Note that the decoding is now purely spatial (accounting for distributed neural representations), while the encoding is purely temporal (capturing only the relevant portions of the stimulus). It is possible to expand both filtering operations to become spatiotemporal, but at the cost of increased dimensionality. The proposed approach is depicted diagrammatically in Figure 7.

**Figure 7:**
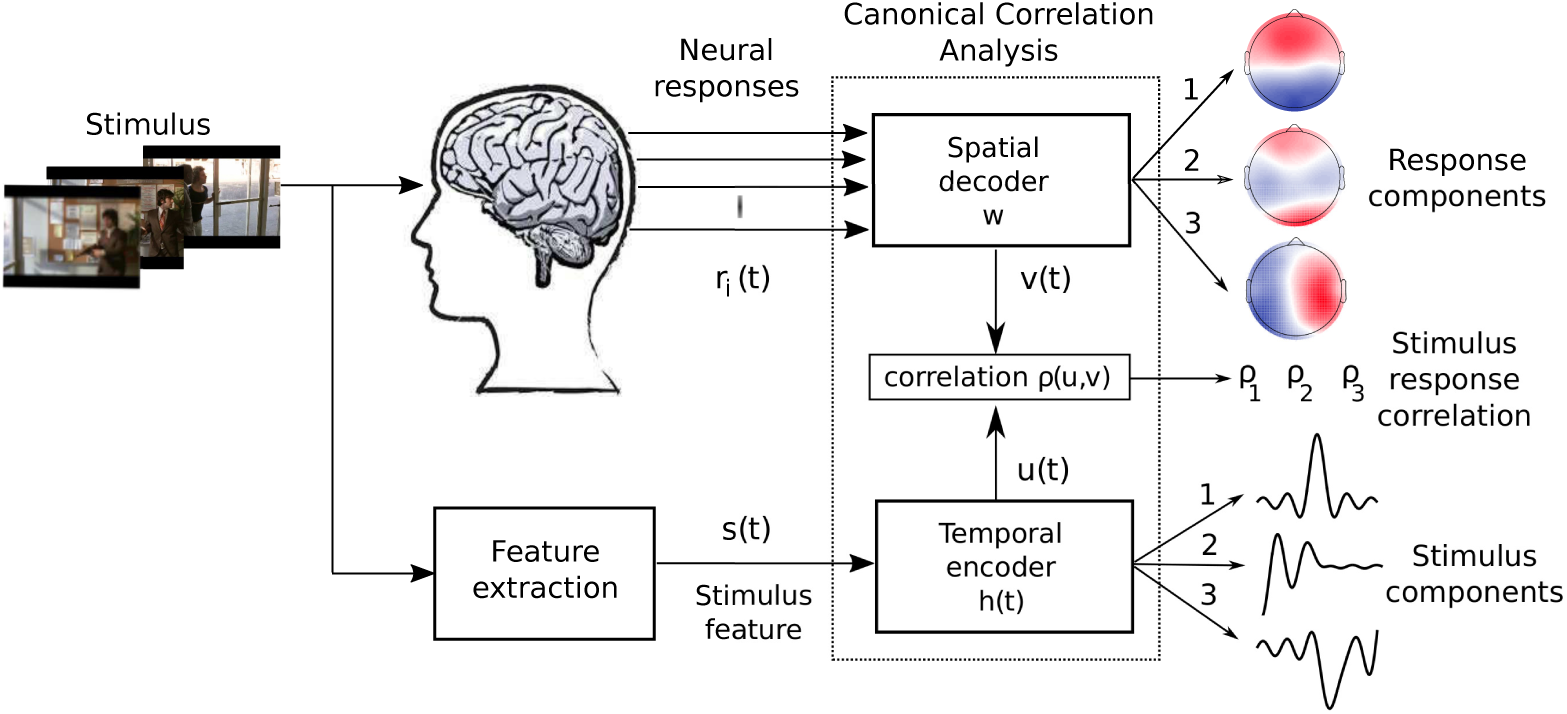
Schematic view of the proposed technique. A stimulus impinges on the observer, generating a spatiotemporal neural response *r*_*i*_(*t*). The relevant features of the stimulus (for example the time-varying luminance or sound envelope) are extracted, resulting in a time series *s*(*t*). An optimization procedure then selects a spatial filter *w* to apply to the neural response and a temporal filter *h*(*t*) to apply to the stimulus such that the resulting filter outputs are maximally correlated in time. The result is a set of multiple response and stimulus components whose activities track each other.

### Comparison and solutions

We summarize the three approaches here so that the analogies can be more clearly identified.

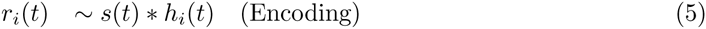

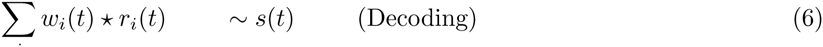

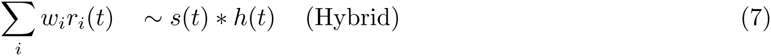

where, *u*(*t*) ~ *v*(*t*) indicates that model parameters are selected to maximize the correlation between the signals *u*(*t*) and *v*(*t*):

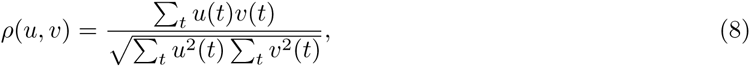

and where zero-mean has been assumed for *u* and *v*. Model parameters that maximize correlation in (5) and (6) are unique and can be found with conventional least-squares optimization, arguably leading to the popularity of the conventional encoding and decoding approaches (see SI 4).

Optimizing the parameters of the hybrid technique (7) is performed by Canonical Correlation Analysis (CCA) [28], which provides *multiple* independent dimensions, or “components” of the correlation between the stimulus and the response. Specifically, each dimension *k* = 1,…,*K* is defined by an encoder *h*_*k*_(*t*) and decoder *w*_*ki*_. Each encoding/decoding dimension *k* captures in 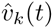 a spatial component of the neural activity and in 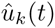 a temporal component of the stimulus. The stimulus-response correlation (SRC) of dimension *k* is then defined according to:

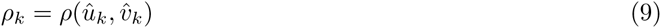

with *ρ*_1_ > *ρ*_2_ >… > *ρ*_*K*_, and zero cross-correlation across components, 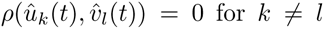 for *k* ≠ *l* (this is approximately true also in the case of regularization demonstrated in the results section). Thus, different components capture genuinely different aspects of the stimulus-response relationship. Note that this multidimensional representation cannot be recovered with the either the encoding approach (5) or the decoding approach (6), as they inherently yield only one dimension of the SRC. Some have used principal components analysis (PCA) as a post-processing of encoding models to capture components of distributed representations [30, 29]. However, PCA enforces orthogonality on the spatial distributions of neural activity. In contrast, CCA only requires temporally uncorrelated components, and spatial filters *w*_*i*_ are not constrained to be orthogonal. CCA has previously been used to establish correlation between stimuli and MEG responses using purely temporal filters [34] without capturing distributed responses.

Note that in the traditional encoding and decoding approaches (5)-(6), the number of free parameters is *QD*, with *Q* representing the length of the temporal aperture and *D* denoting the number of neural data channels. On the other hand, the hybrid method leads to simpler models with (*Q* + *D*)*K* free parameters. Typically, the total correlation is contained within a few components *K* (e.g. 3 or less) while *D* and *Q* often span hundreds of parameters. Thus we expect the new technique to be less susceptible to overfitting the data.

Regularization is routinely performed in order to prevent overfitting, in particular for decoding approaches which simultaneously fit *QD* parameters. Note, however, that regularization is based on the response statistics alone (e.g. by pruning response dimensions with small eigenvalues). In contrast, CCA prunes dimensions based on the criterion of interest (i.e., the SRC) and is thus not at risk of removing relevant dimensions.

### Subjects and stimuli

All participants provided written informed consent in accordance with the procedures approved by the Western Institutional Review Board (Puyallup, WA). 30 healthy human subjects (15 females, median age 23) participated in the main experiment, during which they freely viewed a clip from the film “Dog Day Afternoon” (duration 325s, 24 frames/s; this clip was first analyzed in [25] using fMRI) while their EEG was recorded. An additional 5 healthy human subjects (2 females, median age 21) were recruited for the video game study, where EEG was collected while subjects played the car-racing video game “SuperTuxKart” (variable duration, 60 frames/s). Subjects controlled their vehicle, using their right hand, via keyboard directional arrows -left/right keys controlled steering, while up/down controlled acceleration. The video game study also included a “divided-attention” condition during which participants performed a concurrent rapid-serial-visual-presentation (RSVP) task [15] to earn various “items” which gave players a temporary competitive advantage against the other racers. For this condition, a square black panel was superimposed on the screen with objects rapidly flashed (i.e., 5 Hz) inside the square. Subjects were instructed to attend (while maintaining eye gaze on their vehicle and race track) to a particular object of the RSVP display. Players could redeem their selected items by pressing the space bar with the left hand. Two levels of race difficulty were tested: “easy” and “hard”. The easy condition consisted of slow driving speed against 3 simulated race competitors, tuned to allow the subject ample opportunity to win the race. In the hard condition, the driving speed increased, as well as the number (i.e., 7) and performance of the simulated race competitors. This resulted in more obstacles, crashes, and aggressive driving. In both experiments, sound was delivered via Sony MDR-7506 headphones adjusted by each subject to a comfortable listening volume prior to the experiment.

### Existing data

To evaluate the effect of attention (see Figure 4), we reanalyzed the EEG data from [33]. In that study, EEG was collected during passive viewing/listening of the following popular stimuli: “Bang! You’re Dead” (*N* = 20, 8 females, median age 20, duration 372s, 25 frames/s) [20], “The Good, the Bad, and the Ugly” (same subject pool, duration 388s, 30 frames/s) [21], and “Pie Man” (recorded on a separate *N* = 20 subjects, 7 females, median age 21, duration 360s, 30 frames/s, [40], although this audio narration only showed a fixation cross on the screen). There were two experimental conditions (each with *N* = 20): in the “attend” condition, subjects were instructed to normally attend to the stimuli. To emulate the inattentive state, in the “count” condition subjects were instructed to mentally count backwards in steps of 7 during viewing/listening.

### EEG collection and pre-processing

Subjects were fitted with a 32-electrode cap placed on the scalp according to a modified 10/10 scheme for EEG, which was recorded with a BioSemi ActiveTwo system (BioSemi, Amsterdam, Netherlands) at a sampling frequency of 2048 Hz and 24 bits per sample. Four-channel electrooculogram (EOG) recordings were collected from electrodes below and adjacent to each eye. EEG pre-processing was performed automatically and offline in the MATLAB software (MathWorks, Natick, MA). The signals were high-pass filtered at 1 Hz, notch filtered at 60 Hz, and then downsampled to 256 Hz. To remove the contribution of eye movements from the EEG, the four EOG channels were linearly regressed out of the 32-channel EEG. Artifactual channels and data samples were identified and replaced with zeros when their respective power exceeded the mean power by 4 standard deviations. The EEG was further downsampled to the frame rate of the stimulus prior to analysis.

### Stimulus feature extraction

All stimuli were loaded into the MATLAB software to extract the video frames and audio samples comprising the stimulus. The color video frames were converted to grayscale, resulting in intensity values *I*_*p*_(*t*) for pixel *p* at frame time *t*. The luminance at each frame was then computed as the mean intensity across pixels: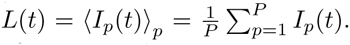 Similarly, temporal contrast was derived as the mean temporal derivative of intensity changes, 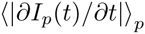 (i.e., unsigned frame to frame changes in intensity). Local contrast was computed following [18]: 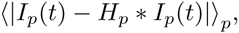 where *∗* indicates a 2-dimensional spatial convolution, here with uniform point-spread function *H*_*p*_ with a 30 × 30 region-of-support. Optical flow was estimated from the frame sequence using the Horn-Schunk method [27] via the MATLAB Computer Vision System Toolbox. The sound envelope was computed as the squared magnitude of the Hilbert transform of the sound pressure amplitude, and then downsampled to the frame rate of the accompanying video. Prior to processing, all features were high-pass filtered at 1 Hz (to match the neural response and remove slow drifts) and z-scored.

### Measuring SRC

SRC were measured for the encoding (5), decoding (6), and hybrid techniques (7). Note that the decoding approach correlates stimulus features, the encoding approach correlates neural responses, while the hybrid approach correlates filtered neural responses with filtered stimulus features. As a benchmark encoding technique, we implemented the “visual evoked spread spectrum analysis” (VESPA) method [38] using the regularization parameter provided therein. The “stimulus reconstruction” decoding technique [53] was also implemented using the publicly available code base from http://www.ine-web.org/software/decoding/, with all parameters set to their default values. All reported correlations were Pearson correlation coefficients cross-validated in a leave-one-out procedure along the subject dimension: model parameters were learned over all but one subject, with the remaining subject serving as the “test-set”. When learning the VESPA model parameters, the channel with the highest correlation on the training set was retained for the test set. Note that all SRC values depend on the spectral content of the signals being correlated. To ensure homogeneity in the various comparisons, we therefore ensured that all stimulus features and neural responses were filtered over a common range (i.e., from 1 Hz to half of the video frame rate). Since not all stimuli had the same frame rate, however, one should be careful in making inferences from differences in SRC between stimuli. In order to detect statistically significant SRC (*ρ* > 0), we formed *N* = 1000 surrogate data records in which the phase spectrum of the EEG was randomized following [64]. This procedure preserved the autocorrelation structure of the EEG while disrupting the temporal relationship between the stimulus and neural activity. SRC computed with the permuted data records defined the null distribution from which p-values were estimated.

### SRC-ISC comparison

When comparing SRCs with ISC (Figure 3), we learned the SRC-maximizing filters (*h* and *w*) separately for each individual subject (i.e., no pooling of subjects’ EEG was conducted for this analysis). This was performed in order to ensure that the computation of SRC-maximizing filters would not be biased towards picking up patterns of activity that were common across subjects, thus confounding the SRC-ISC relationship. For each subject, their SRC-maximizing spatial filter was applied to their EEG record, and the resulting EEG components were then correlated with their corresponding optimally filtered stimulus feature. The correlation was computed in a time-resolved fashion using windows of 5-second length and 80% overlap across successive windows. At each window, the SRC was uniformly summed across the first three components in order to reduce the dimensionality of the comparison. To compute ISCs of the neural responses, we followed the procedure of [12]. The subject-independent spatial filters that maximized ISC across the subject pool were learned and then applied onto each subject’s data before computing pairwise ISCs and then summing across all *N* × (*N* − 1)/2 = 435 subject pairs. As with the SRC, the ISC was computed across 5 second windows and uniformly summed across the first three components.

